# An efficient gene regulatory network inference algorithm for early *Drosophila melanogaster* embryogenesis

**DOI:** 10.1101/213025

**Authors:** Hirotaka Matsumoto, Hisanori Kiryu, Yasuhiro Kojima, Suguru Yaginuma, Itoshi Nikaido

## Abstract

The spatial patterns of gene expression in early *Drosophila melanogaster* embryogenesis have been studied experimentally and theoretically to reveal the molecular basis of morphogenesis. In particular, the gene regulatory network (GRN) of gap genes has been investigated through mathematical modeling and simulation. Although these simulation-based approaches are useful for describing complex dynamics and have revealed several important regulations in spatial patterning, they are computationally intensive because they optimize GRN with iterative simulation. Recently, the advance of experimental technologies is enabling the acquisition of comprehensive spatial expression data, and an efficient algorithm will be necessary to analyze such large-scale data. In this research, we developed an efficient algorithm to infer the GRN based on a linear reaction-diffusion model. First, we qualitatively analyzed the GRNs of gap genes and pair-rule genes based on our algorithm and showed that two mutual repressions are fundamental regulations. Then, we inferred the GRN from gap gene data, and identified asymmetric regulations in addition to the two mutual repressions. We analyzed the effect of these asymmetric regulations on spatial patterns, and showed that they have the potential to adjust peak position. Our algorithm runs in sub-second time, which is significantly smaller than the runtime of simulation-based approaches (between 8 and 160 h, for exmaple). Neverthe-less, our inferred GRN was highly correlated with the simulation-based GRNs. We also analyzed the gap gene network of *Clogmia albipunctata* and showed that different mutual repression regulations might be important in comparison with those of *Drosophila melanogaster*. As our algorithm can infer GRNs efficiently and can be applied to several different network analysis, it will be a valuable approach for analyzing large-scale data.

## 1. Introduction

The investigation of the regulatory mechanisms of pattern formation during development (i.e., morphogenesis) is a fundamental topic in developmental biology. As the molecular basis of morphogenesis, spatiotemporal patterns of gene expression and their regulation have been studied experimentally and theoretically, especially in the context of early embryogenesis in *Drosophila melanogaster* (reviewed in [1, 2]). In the early embryogenesis of *D. melanogaster*, the characteristic spatial expression patterns of gap genes and subsequent patterns of pair-rule genes are established along the anterior–posterior (AP) axis, which is essential for metameric patterning[3]. The gap genes are regulated by maternal effect genes such as *bicoid* and the interactions among gap genes, which results in spatial expression patterns of unimodal or bimodal concentrations along the AP axis (Fig 1(A)). The pair-rule genes are regulated by maternal effect genes and gap genes, and recently, interactions among pair-rule genes have also been suggested to play an important role in producing their striped patterns (Fig 1(B))[4, 5]. Therefore, the elucidation of the interactions among gap genes and pair-rule genes, which entails the inference of the underlying gene regulatory network (GRN), is essential to reveal the mechanisms of spatial expression patterning and morphogenesis.

**Figure 1:**
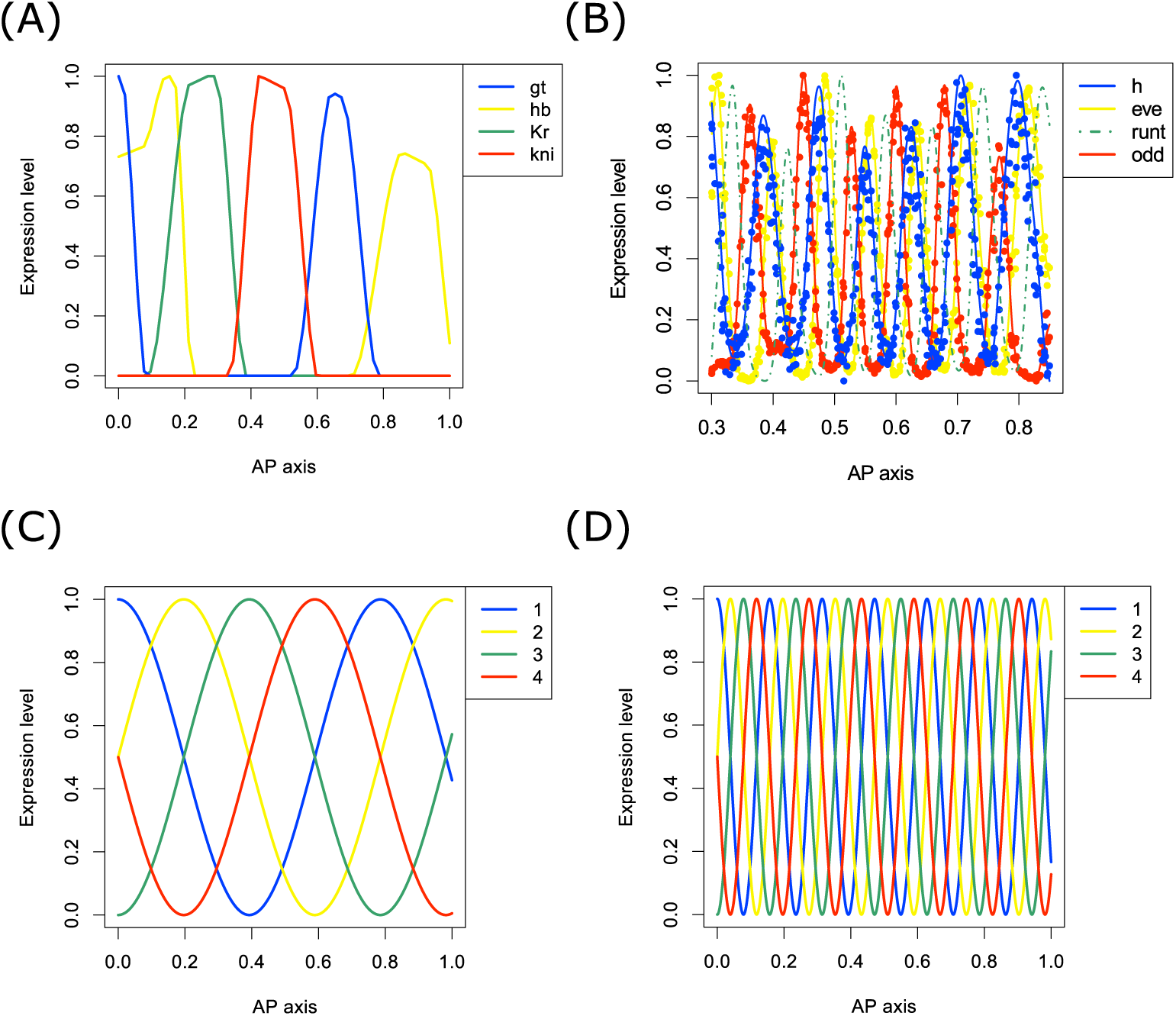
Spatial expression patterns of gap genes and pair-rule genes in *D. melanogaster*. (A) Spatial expression patterns of gap genes (*gt*, *hb*, *Kr*, *kni*) along the anterior–posterior (AP) axis. (B) Spatial expression patterns of pair rule genes (*h*, *eve*, *runt*, *odd*). The filled circles represent the expression values of raw data, and the smoothing curves are drawn for visibility by using the smooth.spline function of R with *df* = 50. Because the expression data for the *runt* gene are not registered in the VirtualEmbryos of the Berkeley Drosophila Transcription Network Project (BDTNP), we represent the *runt* pattern by simply shifting the *eve* pattern (see the Materials and methods section). (C, D) The spatial patterns reproduced by *d***x**(*s*) = **Wx**(*s*)*ds* for *c* = 4 and *c* = 20, respectively.

Several theoretical studies have been conducted to describe the spatial expression patterns of gap genes and infer the GRN by using reaction–diffusion (RD) and similar models[6, 7, 8, 9, 10, 11, 12, 13, 14, 15, 16]. The RD models are difficult to solve analytically, and it is generally difficult to determine parameter values directly from the observed data. Therefore, these researches simulated the model repeatedly while searching for parameter values for which the simulated expression data resemble the observed data. Although such simulation-based approaches have revealed several characteristic properties of spatial patterns and the GRN of gap genes, they are computationally intensive due to repetitive simulation. For example, Jaeger et al. noted that optimization took between 8 and 160 h[6].

As a result of recent advances in sequencing technologies and experimental techniques, the analysis of large-scale spatial expression data has become increasing appealing. For example, a strategy to acquire the spatial transcriptome data has been developed by using positional barcodes [17]. In addition, in situ RNA sequencing[18] and multiplexed RNA imaging[19] will also be useful to acquire spatially resolved expression data. As another example, methods for reconstructing comprehensive spatial expression patterns from single-cell RNA sequencing have been proposed[20, 21]. To fully utilize such large-scale spatial expression data for spatial regulation analysis, an efficient GRN inference algorithm is necessary. Therefore, we have developed an efficient machine learning approach to infer the GRN based on an RD model.

In this approach, we use a linear RD model to describe spatial patterns. We show that the model can be associated with a spatial first-order linear ordinary differential equation (ODE) in the steady-state condition, and the GRN can be derived from the parameters of the ODE, which can be optimized analytically and efficiently by linear regression.

First, we qualitatively analyze our RD model with simulation data, which reproduce the periodic patterns shown in gap genes and pair-rule genes. From this we show that two independent mutual repressions are key regulatory elements; these repressions are also supported by simulation-based research. Then, we apply our algorithm to data for four trunk gap genes (*hunchback* (*hb*), *Kruüppel* (*Kr*), *giant* (*gt*), *knirps* (*kni*)) of *D. melanogaster*. The inferred GRN contains asymmetric repressions, in addition to the two independent mutual repressions. We analyze the influence of these asymmetric regulations and show that an expressed peak position slightly changes in the absence of the regulations, which suggests that these have an effect on the subtle adjustment of spatial patterns. Furthermore, we compare our GRN with two simulation-based GRNs and show that they are highly consistent. The runtime of our algorithm is about 0.25 s, which is significantly smaller than that of simulation-based approaches (between 8 and 160 h, for example[6]).

We also analyze the gap gene network of *Clogmia albipunctata*, and find complete mutual repressions among *hb*, *Kr*, and *knl* (*knl*, which corresponds to *kni* in *D. melanogaster*). Such complete mutual repressions are analytically derived from three-element periodic patterns, and our result suggest that the regulations among *hb*, *Kr*, and *knl* are the fundamental regulations in *C. albipunctata*. Next we evaluate the effect of *gt* based on a model selection approach. Although the analysis suggests that the influence of *gt* is smaller than those of the other three genes, the Akaike information criterion (AIC) values show that *gt* is still important for predicting spatial patterns.

These results show that our algorithm is not only useful for GRN inference but also for other analyses, including model selection, to investigate the GRN evolution. The efficiency of our algorithm will become an increasingly important consideration as the amount and types of data that are available increase.

## 2. Results and Discussion

### 2.1. An efficient GRN inference algorithm based on a linear RD equation

RD equations have been extensively studied as theoretical models for explaining developmental patterning[22, 23, 24]. In particular, several RD based models (or similar models) have been used to describe the spatial patterning of gap genes along the AP axis in early *D. melanogaster* embryogenesis [7, 8, 9, 10, 11, 12, 13, 14, 15, 16]. The general form of the RD equations is

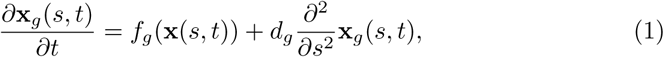

where x(*s, t*) is the expression value vector of four trunk gap genes at time *t* and position *s*. The subscript *g* is an index for the four gap genes. The first term on the right-hand side, *f*_*g*_(x), is a non-linear reaction term for gene *g*, which contains the interaction among gap genes, the regulatory effect of maternal effect genes, and a degradation coefficient. The second term is the diffusion term, and *d*_*g*_ is a diffusion coefficient.

The previous researches have used iterative simulation-based optimization to infer the GRN, which are computationally intensive. In this paper, we use a linear RD based model to efficiently infer the GRN:

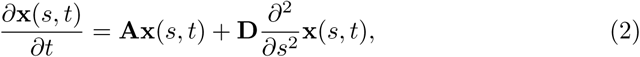

where **A** is a 4 × 4 matrix and **A**_*ij*_ represents the regulatory relation from gene *j* to gene *i*. **D** is a diagonal matrix whose diagonal elements are the diffusion coefficients of each gene.

With the above model, we can derive the following linear second-order ODE under the assumption that the spatial patterns are almost formed and approximately satisfy the steady-state condition (*∂***x**(*s, t*)*/∂t* = 0).

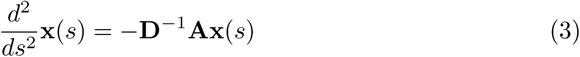

The above second-order ODE can be derived from the following expression and first-order ODE for **x**(*s*):

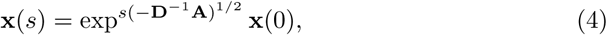

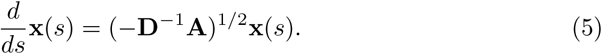

From the above transformation, the linear RD based model can be associated with a spatial first-order linear ODE under the assumption that the system is in a steady state. Therefore, if the spatial patterns can be represented by the spatial ODE *d***x** = **Wx***ds*, then we can derive the GRN in the model (i.e., the matrix **A**) from **W** as follows.

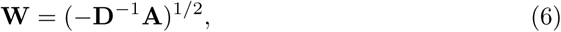

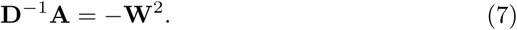

Thus, we can infer the structure of GRN **A** from −**W**^2^ if the diffusion coefficients are almost equal because, in that case, the two matrices are proportional. Thus, in the following, we will refer to −**W**^2^ as the GRN.

From the above equation, we can derive the GRN if we can infer **W** and, therefore, we developed an efficient algorithm to infer **W** from spatial expression data. We approximated the dynamics of *d***x**(*s*) = **Wx**(*s*)*ds* by considering the discretization with a small space step (*δs*):

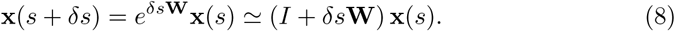

From this approximation, the inference of **W** can be regarded as a linear regression problem if discretized expression data (**x**(*s* + *δs*) and **x**(*s*)) are given:

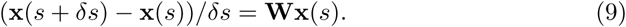

Thus, **W** can be optimized analytically and efficiently by linear regression, and we can derive the GRN easily from −**W**^2^.

### 2.2. Qualitative GRN analysis to validate our algorithm

We first applied our model for simulated data to qualitatively investigate whether the association between our RD based model and the spatial ODE as shown in the previous section is useful for GRN inference. To produce the simulation data, we used a spatial ODE that reproduces the spatial expression patterns of gap genes and pair-rule genes. The spatial patterns of gap genes and pair-rule genes show periodicity along the AP axis such that the four genes in each set are expressed in periodic order: (*gt* -> *hb* -> *Kr* -> *kni*) and (*h* -> *eve* -> *runt* -> *odd*) (see Fig 1(A,B)). We used the following spatial ODE to reproduce these periodic patterns along the AP axis.

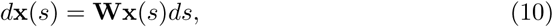

where **x**(*s*) is a 4-element expression vector at position *s*, and **W** is a 4×4 matrix that reproduces periodic patterns: **W**_*i,j*_ = −**W**_*j,i*_ = *c* when (*i, j*) is (2, 1), (3, 2), (4, 3), or (1, 4), and **W**_*i,j*_ = 0 otherwise (see Fig 2(A)). Fig 1(C,D) show the spatial patterns based on the above ODE with *c* = 4, 20, respectively. The results recover the periodicity shown in the gap and pair-rule gene expression patterns.

**Figure 2:**
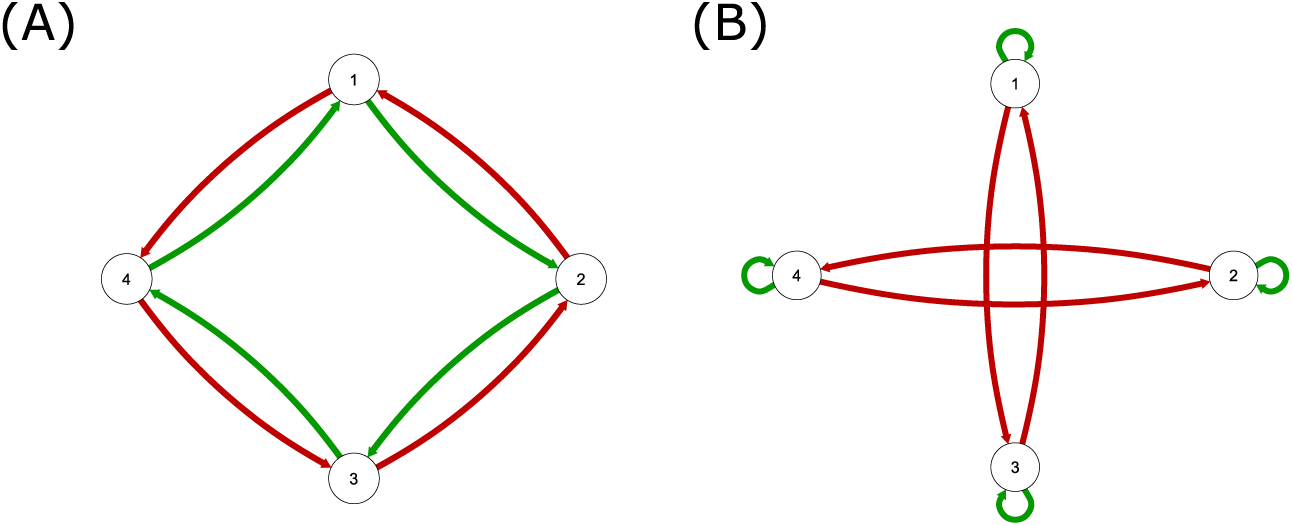
Graphical representations of **W** and −**W**^2^. (A) The network structure of **W**. (B) The network structure of −**W**^2^, which corresponds to the gene regulatory network. The green and red arrows represent positive and negative regulatory relations, respectively.

Then, we derived the GRN in our model (−**W**^2^) by using the matrix **W** (Fig 2(B)) given above. In the resulting GRN, two mutual-repression regulatory relations are represented. Given the correspondence of the gap genes (or pair-rule genes) to the patterns generated by *d***x**(*s*) = **Wx**(*s*)*ds*, the mutually repressing gap gene pairs are (*gt*, *Kr*) and (*hb*, *kni*), and (*h*, *runt*), and (*eve*, *odd*) are the mutually repressing pair-rule genes. These mutually repressing regulations have been noted as being important for gap genes[25] and pair-rule genes[4]. In particular, the two independent mutual repression regulations, which are referred to as the two parallel toggle switches, are considered to be the critical components of the gap gene network[26]. In addition, mutual repression has been suggested to be important from the perspective of precise and stable boundary formation in spatial expression patterns[27, 28]. Although the French flag model[29], which describes spatial patterning solely from the gradient of expression of genes, such as maternal effect genes, can reproduce global spatial patterns, theoretical studies suggest that accurate local boundary formation is difficult to reproduce from this model alone. However, this problem can be solved by including the RD model and mutual repression in the French flag model [28]. Thus, the approximated GRN −**W**^2^ comprises important regulations in the networks of gap genes and pair-rule genes, and our RD based model will be useful for GRN inference.

Next, we investigated whether the GRN can be inferred from spatial data using our optimization algorithm. We constructed spatially discretized data from the spatial ODE with *c* = 4 and optimized 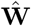. Fig 3(A) shows the spatial patterns reconstructed with 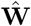, and Fig 3(B) shows the comparison between the true GRN (−**W**^2^) and the inferred GRN 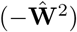. The Pearson’s correlation coefficients between the reconstructed spatial patterns and the true patterns, and between the inferred GRN and the true GRN are over 0.99. Thus, our algorithm can successfully infer **W** and the GRN from spatially discretized data.

**Figure 3:**
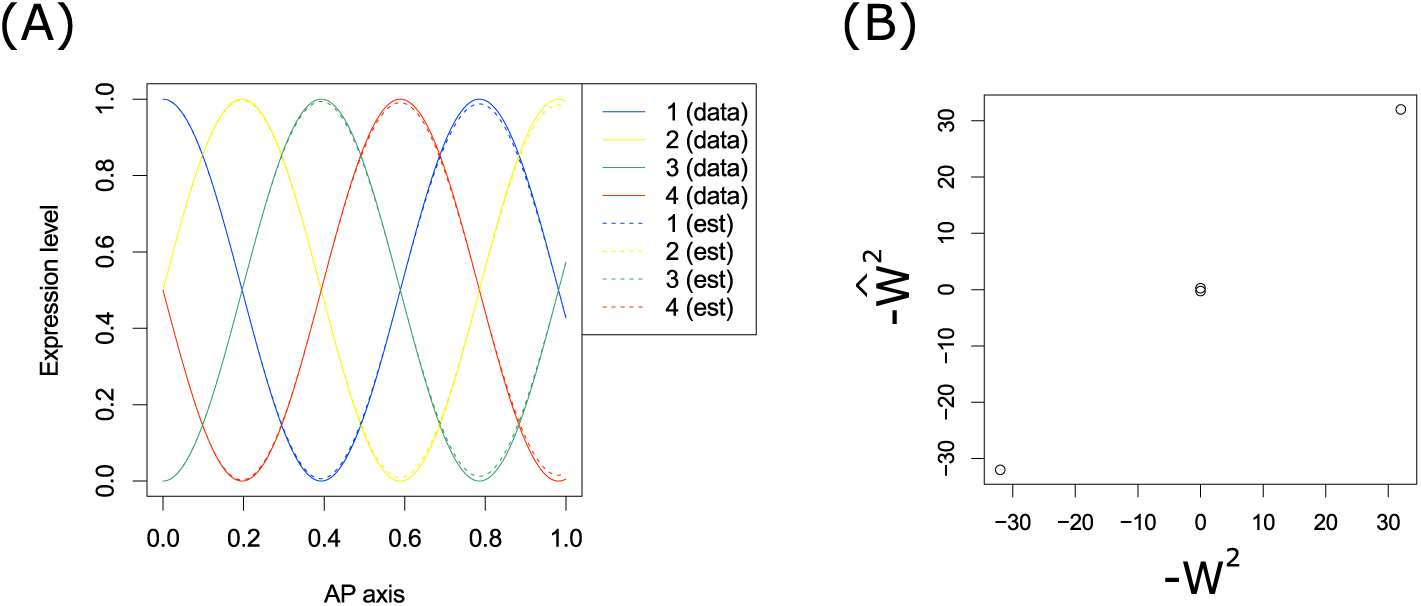
Evaluation of optimized 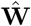 for simulation data. (A) The spatial patterns of true data (full lines) and reconstructed patterns from 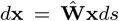 (dotted lines). (B) Comparison between the true GRN (−**W**^2^) and the estimated GRN 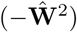.

### 2.3. Inference of gap gene network of Drosophila melanogaster

We have evaluated the GRN derived from virtual spatial patterns thus far. Although the simulation data are useful for qualitative analysis, it does not completely represent the observed expression patterns and, therefore, the above GRN may not contain all the regulatory relationships. In this section, we infer the GRN from the real gap gene expression patterns.

We first calculated the optimized parameter 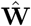 from the observed gap patterns with our algorithm. We evaluated the linear regression models for predicting each of the four genes based on the p-values of F-statistics. The values are significantly small: the values of − ln(p-value) are about 235, 184, 229, and 69 for *hb*, *Kr*, *gt*, and *kni*, respectively. To validate the optimization for total spatial patterns, we reconstructed the patterns with 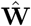 (Fig 4(A)). Because of the limits of a linear model, the optimized model cannot represent the sharp peaks shown in the observed data. However, the positions of peaks are almost the same in the data and the reconstructed patterns; therefore, we concluded that our model discovered the fundamental expression patterns (Fig 4(C)).

**Figure 4:**
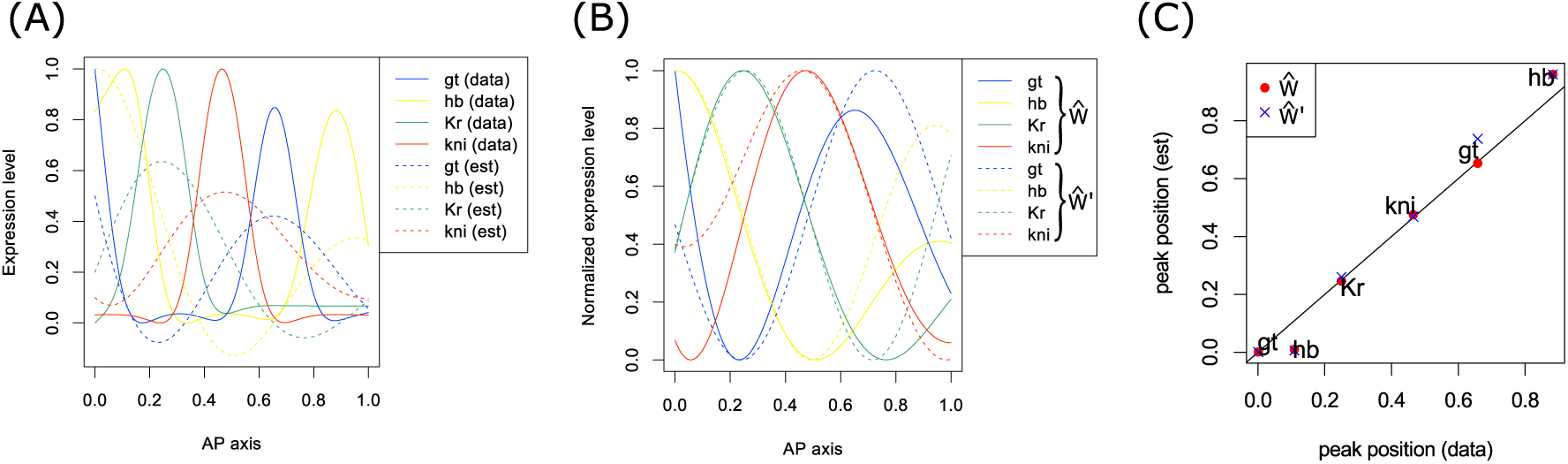
Evaluation of optimized 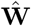 for gap gene data. (A) The spatial patterns of input data (gap gene data after smoothing) (full lines) and reconstructed patterns from 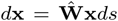 (dotted lines). (B) The reconstructed patterns from 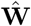 (full lines) and those from 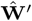 (dotted lines). (C) Comparison of the peak positions for input data and reconstructed data. The filled circles and crosses correspond to the patterns of 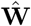 and 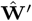, respectively.

Next, we derived the GRN (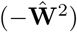) from the above 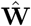 (Fig 5(A)). In the GRN inferred from virtual expression data, the regulatory edges corresponding to the mutually repressing regulations of (*gt*, *Kr*) and (*hb*, *kni*) have large negative values. In addition, the edges from *hb* to *gt* and from *Kr* to *hb* have large negative values, while the edges from *gt* to *kni* and from *kni* to *Kr* have small negative values. Such repressions are supported by previous research and are described as weak asymmetric repressions between overlapping gap genes[25]. To evaluate the effects of such asymmetric regulations, we reconstructed the spatial patterns with 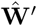 so that 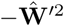 only has two mutual repressions (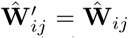 for (*i, j*) corresponding to (*gt*, *hb*), (*hb*, *gt*), (*hb*, *Kr*), (*Kr*, *hb*), (*Kr*, *kni*), (*kni*, *Kr*), (*kni*, *gt*), and (*gt*, *kni*), and 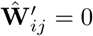 otherwise) and compared these patterns with the patterns reconstructed from 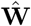 ((Fig 4(B)). Although most peak positions are almost equal, the positions of the posterior peak of *gt* are slightly different. The actual posterior peak position of *gt* is almost equal to the peak position derived from 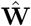 (Fig 4(C)). Therefore, the regulations estimated in addition to the two mutually repressing regulatory interactions might cause a subtle adjustment of spatial patterns.

**Figure 5:**
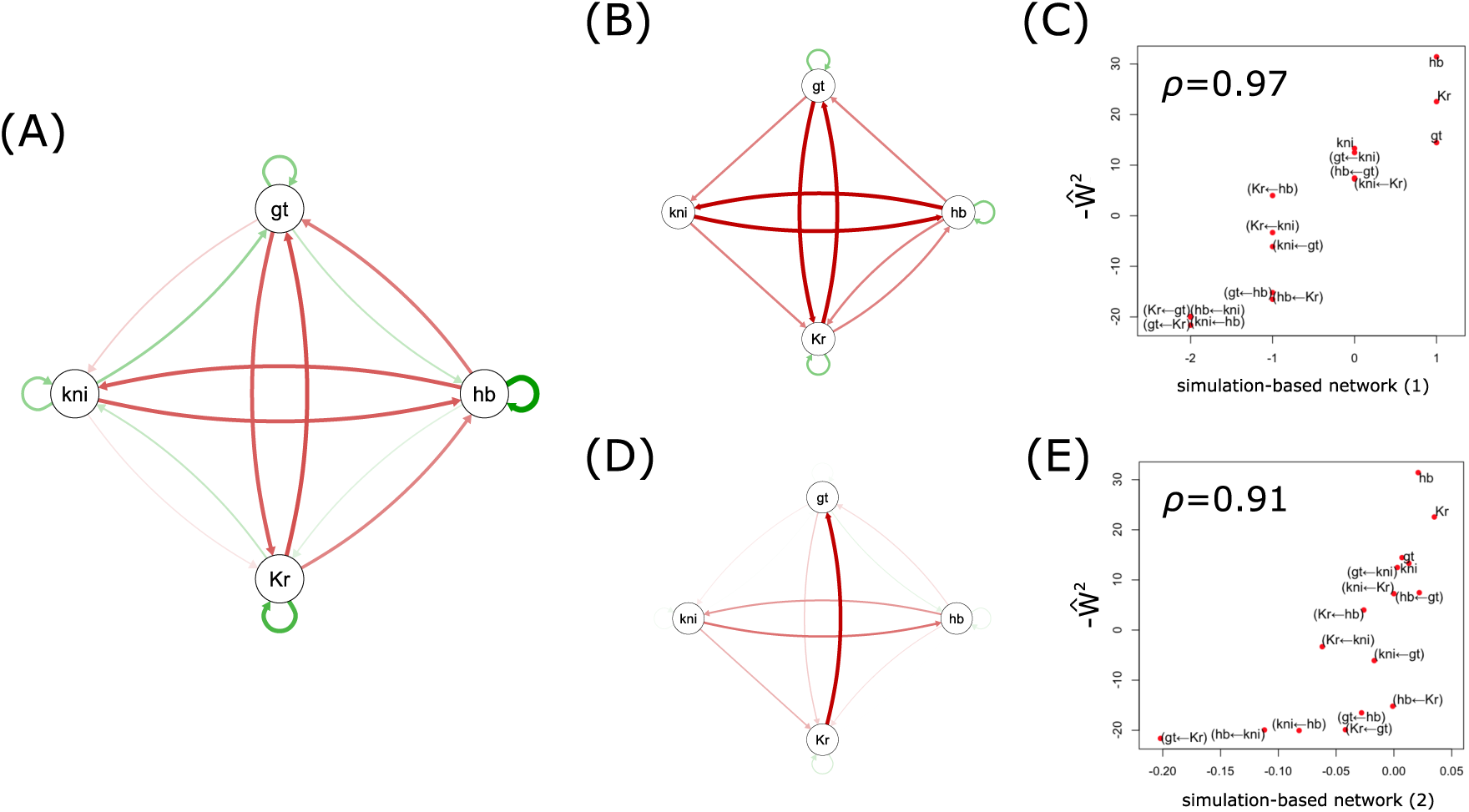
Inferred gap gene networks. (A, B, D) The gap gene networks inferred by our algorithm, Perkins et al.[10], and Manu et al.[13], respectively. Red and green arrows represent negative and positive regulatory relations, respectively, while the thickness of the arrow represents the strength of the relation. Because the GRN of Perkins et al. is a qualitative network, we classified the repressions into strong repression (-2) and weak repression (-1) based on previous research[25]. The GRN of Manu et al. is determined from the values *T*^*a←b*^ where *a* and *b* represent the gap gene indexes, as described in the Supplementary Table of [13]. (C, E) Comparison between our GRN and the GRNs of Perkins et al. and Manu et al., respectively. The filled circles show the data points and the proximal text indicates the regulation correspond to the data point.

To validate our inferred GRN (Fig 5(A)) we compared it with two GRNs optimized using a simulation-based approach (Fig 5(B,D)). The first simulation-based GRN is derived from the combined model inferred by Perkins et al.[10] (Fig 5(B)). The simulated GRN is comprises only qualitative regulatory relationships (activation and repression) so we classified the repressions into strong and weak repressions based on previous research[25]. The Spearman’s rank correlation coefficient between our GRN and the simulation-based GRN is about *ρ* = 0.97 (*ρ* = 0.94 for non-diagonal elements)(Fig 5(C)). The second simulation-based GRN is that inferred by Manu et al.[13] (Fig 5(D)). The Spearman’s rank correlation coefficient between our GRN and the simulation-based GRN is about *ρ* = 0.91 (*ρ* = 0.87 for non-diagonal elements)(Fig 5(E)). Therefore, our model, which assumes a linear reaction and a steady state, can infer a network that is highly consistent with those obtained by more computationally intensive simulation methods. We also used the diffusion coefficients estimated by Manu et al.[13] to investigate the influence of **D** on GRN inference. The Pearson’s correlation coefficient between 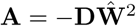 and 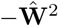 was 0.97 and, therefore, we concluded that the network structure can indeed be approximated by −**W**^2^.

Because our algorithm simply optimizes **W** using linear regression, the run times are significantly shorter than those of other approaches. In the inference of the gap gene network, the runtime of our algorithm was about 0.25 s, which is significantly smaller than those of simulation-based algorithms (for example, Jaeger et al. noted that optimization took between 8 and 160 h[6]). Thus, our algorithm is efficient and, therefore, will be very useful, especially when the number of genes in the GRN increases.

### 2.4. Inference of gap gene network of Clogmia albipunctata

We have investigated the GRN of *D. melanogaster* thus far. In this section, we consider the GRN of *C. albipunctata*. Fig 6 shows the gap gene expression patterns of *D. melanogaster*, *Megaselia abdita*, and *C. albipunctata*. Although the expression patterns of *D. melanogaster* and *M. abdita* are similar, *C. albipunctata* shows a distinctly different spatial pattern. In *C. albipunctata*, *knl*, which corresponds to *kni* in *D. melanogaster*, has a bimodal expression pattern. To investigate such differences from the perspective of the GRN, we optimized **W** from the patterns for *C. albipunctata* (the values of − ln(p-value) based on F-statistics of regression models are about 1340, 1597, 289, and 2043 for *hb*, *Kr*, *gt*, and *knl*, respectively), reconstructed the patterns (Fig 7(A)), inferred the GRN (Fig 7(B)), and compared the GRN of *C. albipunctata* with that of *D. melanogaster* (Fig 7(C)). Two mutual repression pairs (*hb*, *Kr*) and (*Kr*, *knl*) were detected, in addition to the pairs (*gt*, *Kr*) and (*hb*, *knl*). These mutual repressions, which were not detected in the GRN of *D. melanogaster*, construct a complete mutual repression network among *hb*, *Kr*, and *knl*. In previous simulation-based research on *C. albipunctata*, such mutual repression interactions among these three genes have been partially suggested[30]. Such interactions among three genes can be analytically derived from a three-dimensional periodic simulation model (**W**_1,2_ = **W**_2,3_ = **W**_3,1_ = *−c*, **W**_2,1_ = **W**_3,2_ = **W**_1,3_ = *c*) ((Fig 7(D,E))). Here, if we exclude *gt*, the three remaining gap genes may be regarded as having a periodic pattern (i.e., *knl* -> *hb* -> *Kr*) in *C. albipunctata*. Therefore, in contrast to the two independent mutual repressions in *D. melanogaster*, the three gene (*hb*, *Kr*, and *knl*) regulations might be the main mechanisms sustaining the spatial pattern of gap genes in *C. albipunctata*.

**Figure 6:**
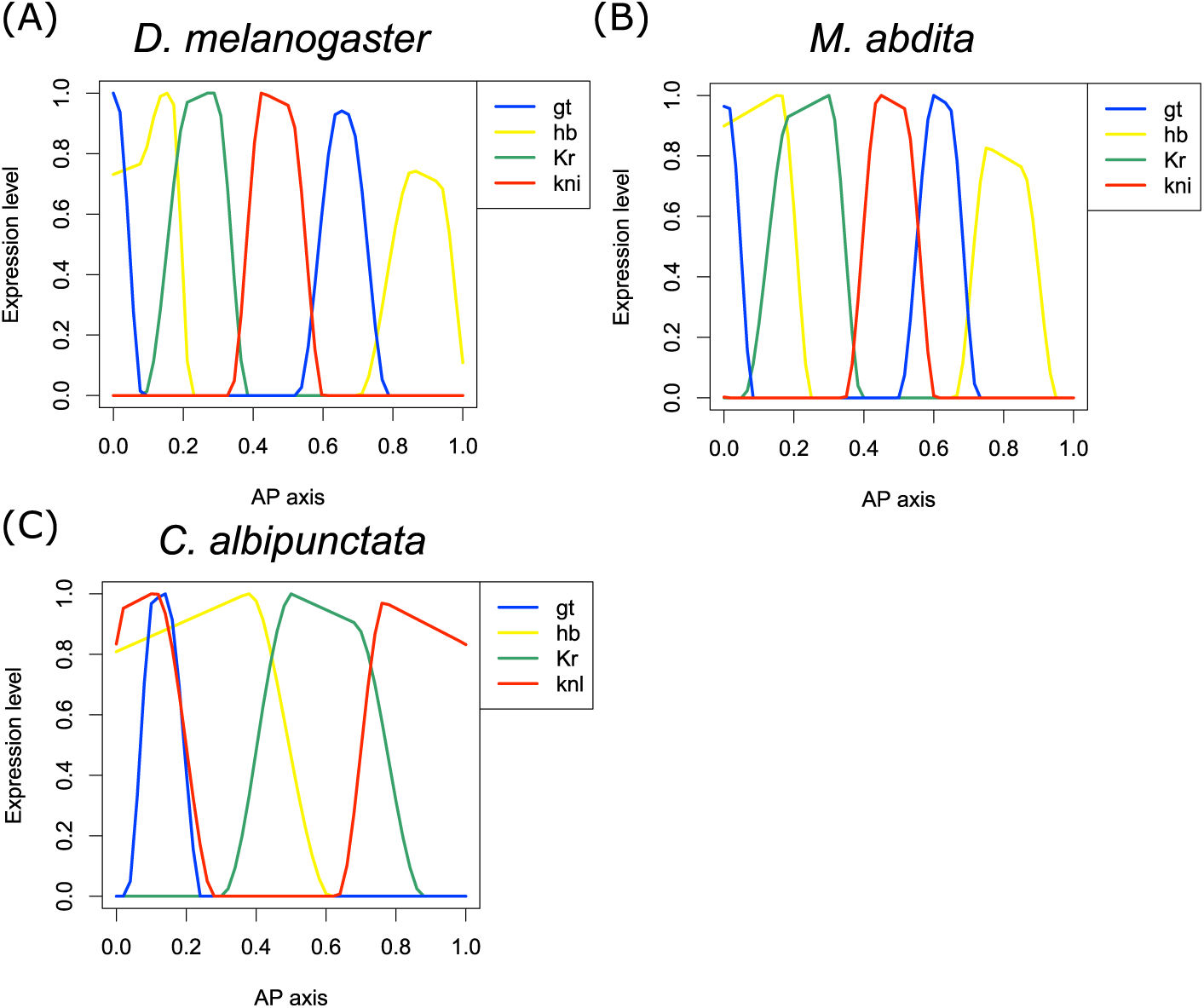
**Spatial expression patterns of gap genes from** *Drosophila melanogaster* **(A),** *Megaselia abdita* **(B), and** *Clogmia albipunctata* **(C).**

**Figure 7:**
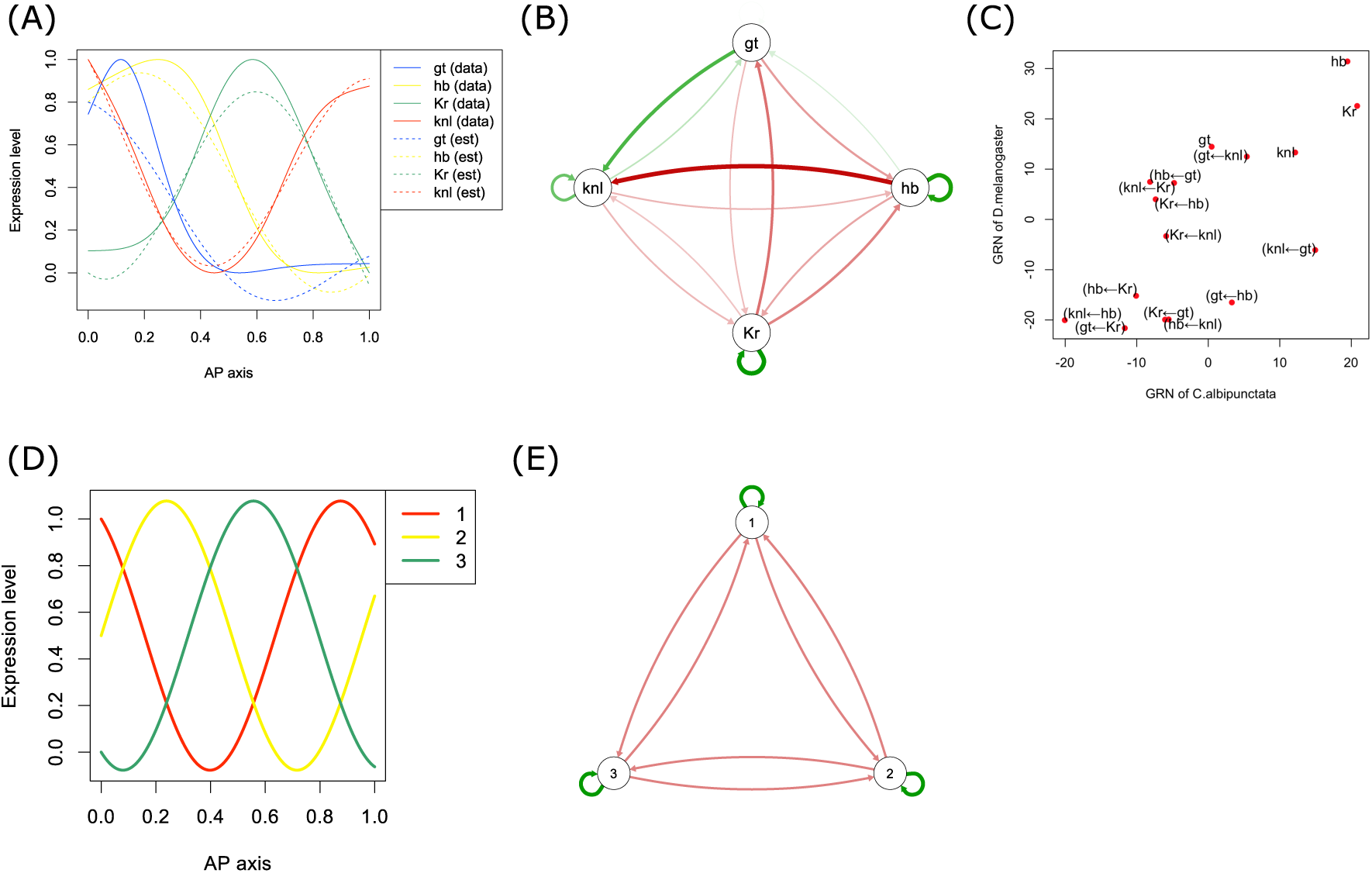
**Analysis for** *C. albipunctata*. (A) The spatial patterns of smoothed input data for *C. albipunctata* (full lines) and patterns reconstructed from 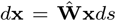 (dotted lines). (B) Estimated gap gene network of *C. albipunctata*. (C) Comparison between the GRN of *C. albipunctata* and that of *D. melanogaster*. (D,E) The simulation data which reproduce the three periodic patterns (D) and the GRN derived from the simulation data (E).

Next, we investigated whether *gt* is necessary for predicting the spatial patterns based on the model selection approach. We compared the AIC value of a 3-gene model, in which one gene is excluded from the explanatory variables, with that of a full model (see Materials and methods section for the detailed procedure). The differences of the AIC values (AIC^∆*g*^(*i*) − AIC^*all*^(*i*)) for each 3 gene model (∆*g*) for predicting the pattern of gene *i* are shown in Table 1. The average for the ∆*gt* model is smaller than those of the ∆*hb*, ∆*Kr*, and ∆*knl* models and, therefore, the effect of *gt* will be smaller than that of the other three genes. However, the values AIC^∆*gt*^(*i*) − AIC^*all*^(*i*) are positive, which means that the pattern of *gt* is necessary for predicting the expression patterns of other genes and, therefore, the regulations from *gt* will also be important in the gap gene network of *C. albipunctata*.

**Table 1:** **Effects of each gene for predicting spatial patterns of each gene.**

|  | *gt* | *hb* | *Kr* | *knl* | average |
| --- | --- | --- | --- | --- | --- |
| $\Delta$ *gt* | | 399 | 502 | 31 | 311 |
| $\Delta$ *hb* | 330 | | 2004 | 2202 | 1512 |
| $\Delta$ *Kr* | 41 | 1463 | | 2806 | 1436 |
| $\Delta$ *knl* | 2 | 395 | 1739 | | 712 |

The row headers represent the excluded gene and the column headers represent the target gene for prediction (the last column gives the average). The values are the corresponding $AIC^{\Delta g}(i) - AIC^{all}(i)$ , where $g$ is the excluded gene (row) and $i$ is the target gene (column).

The results from our analysis of the GRN for *C. albipunctata* show that our algorithm will be useful not only for GRN inference of a single species but also for comparing GRNs among different species. Moreover, our algorithm can be easily applied to model selection, and can evaluate the effects of genes. Therefore, our algorithm is flexible as well as efficient, and it will be useful for further GRN analyses.

In addition to the above analyses, the properties of our GRNs in which the mutual repressions are fundamental regulations can be used to discuss the contribution of GRN evolution to phenotypic evolution. The spatial expression patterning of such pair-rule genes can be classified into two types in arthropods: short-germ and long-germ patterning. The short-germ type, which has been suggested to be the ancestral type, forms striped expression patterns sequentially. Sequential patterning has been suggested to be formed by oscillational expression at the ends of the axis and axis elongation, which are the mechanisms of the clock and wavefront models[31]. In the short-germ type, negative feedback is detected [32, 33], which may be a key regulatory interaction causing oscillations[31]. In contrast, the long-germ type forms spatial patterns almost simultaneously. Under such simultaneous patterning, the importance of the mechanisms forming accurate boundaries increases, and mutual repression contributes to these mechanisms. Therefore, the evolution of mutual repression might be a key factor in the evolution of the long-germ patterning phenotype.

## 3. Materials and methods

### 3.1. Dataset for gap gene expression

We used processed expression data for four trunk gap genes of *D. melanogaster*, expressed along the AP axis and available from SuperFly (http://superfly.crg.eu)[34]. SuperFly contains spatial expression data at several time points throughout early embryogenesis, and final time point data were used in this study. The expression data for each gene were normalized so that the maximum value was equal to 1.0, and the spatial expression patterns are shown in Fig 1(A).

We also generated short-interval spatial expression data from the raw dataset, which consists of expression data for 53 points in space, to estimate **W** via linear regression. First, we rescaled the position variables from 0.0 to 1.0 by setting the offset to 0.0 and the interval length to 1/52. Next, we estimated the smoothing function to interpolate the raw dataset by using the smooth.spline function of R (the parameter *df* is set to 10). Then, we reconstructed the expression data whose interval length is 0.001 by using the above smoothing function, which resulted in a matrix consisting of four genes each with 1,001 position expression values. Last, based on the above reconstructed expression matrix, we generated the target variables (**x**(*s* + *δs*) − **x**(*s*))*/δs* with *δs* = 0.001 and explanatory variables **x**(*s*) for *s* = 0, 0.001*, …,* 0.999, which resulted in 1,000 datasets for each gene.

We also used the gap gene expression data of *M. abdita* and *C. albipunctata* as recorded in SuperFly. Because the *C. albipunctata* data contained the expression values from end to end along the AP axis, consisting of 100 points in space, we removed data from both ends and used the data from points 30 to 80. The spatial patterns of *C. albipunctata* are more gentle than those of *D. melanogaster*, and we used *df* = 5 for estimating the smoothing function. Then, we generated the dataset for linear regression in the same way described for processing the *D. melanogaster* data.

### 3.2. Dataset for pair-rule gene expression

We used the VirtualEmbryos in Expression Atlases dataset downloaded from the BDTNP (http://bdtnp.lbl.gov/Fly-Net/bidatlas.jsp)[35] to obtain expression data for the pair-rule genes (*h*, *eve*, and *odd*) of *D. melanogaster*. To extract the expression data along the AP axis, we used data from the central dorsal–ventral (DV) regions (data within the −10 to 10 DV position range in VirtualEmbryos). We also extracted the data from the 30% to 85% AP-axis region to focus on the region in which the spatially periodic expression patterns appear. Next, we normalized the above data so that the maximum and minimum values were equal to 1.0 and 0.0, respectively, for each gene. The resulting spatial patterns are shown in Fig 1(B). Because VirtualEmbryo does not contain expression data for the gene *runt*, we represented the spatial pattern of *runt* by shifting the expression pattern of *eve* so that the peaks formed the periodic pattern of the four genes (i.e., *h* -> *eve* -> *runt* -> *odd*) found in previous research[36].

### 3.3. Parameter optimization and model selection

We optimized **W** from the 4×1000 target variable matrix **Y** and the 4×1000 explanatory variable matrix **X**. **Y**_*i,j*_ and **X**_*i,j*_ are the *i*-th elements of (**x**(*s* + *δs*) − **x**(*s*))*/δs* and **x**(*s*), respectively, where *s* is the position corresponding to the index *j* (*s* = 0.001 * (*j* − 1) and *δs* = 0.001 in this case). Then, **W** can be optimized as follows.

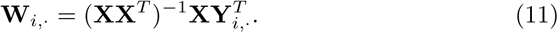

In the case that the inverse of **XX**^*T*^ cannot be computed due to collinearity (e.g., for the simulation data in the qualitative analysis section), we used the Moore–Penrose generalized inverse instead. For spatial pattern reconstruction from the optimized **W**, we used grid search to define the initial value **x**(0) so that the residual sum of squares between the reconstructed patterns and **x**(*s*) is smallest.

We evaluate the effect of a gene using the model selection approach. First, we calculated the AIC value for predicting target variables of a gene *i* (**Y**_*i,·*_) with **W**_*i*,·_**X** (AIC^*all*^(*i*)) by using the lm and AIC functions of R. We refer to this AIC as AIC^*all*^(*i*). Next, we calculated the AIC of the model that excludes gene *g* to predict the same gene *i*, **W**_*i*,·_**X**_*−g,·*_ and refer to this AIC as AIC^∆*g*^(*i*). Then, we calculated the difference of AIC values between each 3 gene model and the full model for each target gene (AIC^∆*g*^(*i*) − AIC^*all*^(*i*)).

## 4. Conclusion

In this study, we developed an efficient GRN inference algorithm to investigate the regulatory mechanisms underlying the formation of spatial expression patterns of gap genes and pair-rule genes in early embryogenesis.

We first qualitatively investigated the properties of the GRN with simulation data and identified that two mutual repressions are the fundamental regulations in spatial periodic patterns. Such mutual repressions have been theoretically suggested to be important for accurate boundary formation. Therefore, these regulations will be fundamental regulations in the gap gene and pair-rule networks. Then, we inferred the gap gene network of *D. melanogaster* and identified asymmetric regulations in addition to the mutual repressions. We investigated the effect of such regulations and showed that they might have a subtle effect on the spatial patterns.

We also compared our GRN with the GRNs inferred from simulation-based approaches, showing that our GRN is highly consistent with these GRNs. However, the runtime of our algorithm is less than 1 s, which is significantly smaller than those of simulation-based approaches which typically take between 8 and 160 h. Recently, comprehensive spatial expression data have been obtained, and our efficient algorithm will be useful to analyze such large-scale data.

Finally, we investigated the GRN of *C. albipunctata* and identified mutual regulations that are different from those of *D. melanogaster*. We also analyzed the effect of gap genes in pattern prediction based on a model selection approach. These results show that our algorithm is efficient and flexible enough to be used not just for spatial patterning analysis but for several other types of analysis, such as GRN evolution.

## 5. Acknowledgments

The authors thank the members of our laboratory for assistance in this study. This work was supported by the Japan Science and Technology Agency (JST), a Grant-in-Aid for Japan Society for the Promotion of Science (JSPS) Fellows, and JSPS KAKENHI Grant Number 16J05079.

